# The Evolution of Early-Life Effects on Social Behaviour – Why Should Social Adversity Carry Over to the Future?

**DOI:** 10.1101/423764

**Authors:** Bram Kuijper, Rufus A. Johnstone

## Abstract

Numerous studies have shown that social adversity in early life can have long-lasting negative consequences for social behaviour in adulthood, consequences that may in turn be propagated to future generations. Given these intergenerational effects, it is puzzling why natural selection might favour such sensitivity to an individual’s early social environment. To address this question, we model the evolution of social sensitivity in the development of in helping behaviours, showing that natural selection indeed favours individuals whose tendency to help others is dependent on early-life social experience. We find that natural selection typically favours positive social feedbacks, in which individuals who received more help in early life are also more likely to help others in adulthood, while individuals who received no early-life help develop low tendencies to helping others later in life. This positive social sensitivity is favoured because of an intergenerational relatedness feedback: patches with many helpers tend to be more productive, leading to higher relatedness within the local group, which in turn favours higher levels of help in the next generation.

## 1 Introduction

In many taxa, the social environment experienced during early life gives rise to predictable between-individual differences in adult social behaviour [1–5]. For example, in many rodents, individuals who have received limited parental care also provide less parental care themselves to their own offspring [6, 7]. Similarly, macaques exposed to parental abuse are likely to develop similar abusive traits in adulthood [8]. These examples hint at a form of developmental plasticity, in which social cues early in life lead to irreversible developmental switching among different social phenotypes [3, 5, 9]. A commonly found pattern is that early-life social adversity results in the reduced expression of prosocial behaviour later in life, which in turn results in social adversity in the next generation [7, 10, 11].

The long-term negative consequences of early-life social adversity, which may even spill over into the next generation, raise the question of why sensitivity to early social experiences has evolved at all: if anything, one might expect that offspring are selectively favoured to buffer the effects of early life adversity, thus preventing the transmission of adverse social behaviours to future generations [12], yet this does not seem to occur. To understand why developmental plasticity is favoured by natural selection, a growing body of theoretical work therefore suggests that early life effects may be an adaptive response to information about potential future environments (e.g., 13–21), suggesting that social adversity during early life is indicative of future social adversity, thus favouring the development of a less social phenotype. While this explanation is intuitive, a key shortcoming of these models is that they have been exclusively formulated with abiotic environments in mind, whereas in the context of social behaviours, the future is shaped by the actions of individuals themselves. Consequently, these models cannot explain why individuals who have experienced social adversity early in life are selectively favoured to go on and subsequently create a socially adverse environment for their own offspring [3, 22].

To understand how early-life social experiences can lead to the intergenerational transmission of socially benign or adverse conditions, we develop an evolutionary model of a developmentally plastic social trait. We focus on the evolution of a helping in a patch-structured population, in which individuals make an irreversible decision early in life to develop either as a nonreproductive helper or as a potential reproductive adult; an individual’s strategy determines the probability with which it develops as a helper rather than a breeder (e.g., [23–26]). The exact number of helpers recruited to a patch varies, with the expected number proportional to the average helping tendency expressed by local individuals. In line with the majority of the theoretical literature, we assume that helping is assumed to increase the fecundity of the reproductives in the local group [27]. The helping tendency expressed by a newborn can then evolve to become dependent on the number of helpers in the local patch present at the time of birth, reflecting the results of empirical studies in which developmental plasticity is based on the current social structure of the local group (e.g., [4, 28, 29]). We then study whether social behaviours are indeed likely to become sensitive to social experience in early life, and if so, what form such developmental plasticity takes.

## 2 The Model

We consider a demographically explicit model of a sexually reproducing metapopulation with non-overlapping generations, distributed over infinitely many demes (Wright’s infinite island model [24, 30]). Each deme contains *n*_b_ adult breeders, who are assumed to reproduce as simultaneous hermaphrodites for the sake of tractability. In addition to breeders, demes can also contain *j* nonreproductive helping individuals, thereby positively affect the fecundity of their reproductive patch mates. We assume that individual demes vary in the number 0 ≤ *j* ≤ *n*_h,max_ of helpers that have successfully been recruited (see the paragraph “Life cycle” below). To assess whether social experiences in early life effect later-life helping, we then ask whether the decision of newborns to become helpers evolves to be dependent (i.e., conditional helping, [31]) on the number of helpers currently present on the patch. Below, we provide a verbal summary of the life cycle, while an extensive description is given in Section 1 of the Online Supplement.

### 2.1 Life cycle

Consider a focal mutant adult breeder who lives on a patch with *n*_b_ − 1 other breeders and *j* nonreproductive helpers. It randomly chooses a mate from among the *n_b_* breeders in the local patch and subsequently produces a large number of *f*(*j*) offspring. Here, fecundity *f*(*j*) is an increasing function of total amount of help received from the *j* helpers, which we assume to be equally distributed over the *n*_b_ breeders present in the patch. A juvenile born from the mutant focal breeder will forego on reproduction and develop as a helper with probability *h*^•^ (*j*), where ^•^ indicates the helping tendency expressed by the mutant mother (differing slightly from the average helping tendency *h*(*j*) in the population). As mentioned before, a key assumption of our analysis is that the tendency to develop as a helper can evolve to become dependent on the current number of helpers in the local patch *j*. Alternatively, with probability 1 − *h*^•^ (*j*), a juvenile does not develop as a helper, in which case it either disperses to a randomly chosen remote patch with probability *d* or remains at the local patch with probability 1 − *d*.

After dispersal, all non-helping juveniles, both philopatric and immigrant, then compete for the *n*_b_ breeding positions. The cycle then repeats, with the newly established breeders’ fecundity now affected by a number of *k* helpers, recruited from the helping juveniles born in the local patch. Specifically, we assume that the number of helpers in each patch is given by a truncated Poisson distribution, where the probability *s*_*j*→*k*_(*h_j_*) that a local patch which previously contained *j* helpers now contains *k* helpers is given by

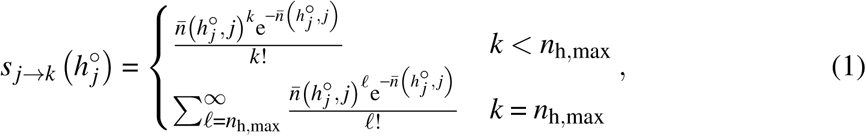

where the first line reflects the Poisson probability of sampling *k* helpers when 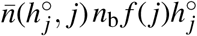 is the average number of helping juveniles produced by all adult breeders in the local patch. The second line reflects the probability that the maximum of *k* = *n*_h,max_ helpers is attained, which occurs when *ℓ* = *n*_h,max_ helpers are sampled, or when more helpers than positions available for them are sampled (i.e., *n*_h,max_ < *ℓ* < ∞), in which case we assume that helpers compete amongst themselves for the *n*_h,max_ available helping positions, with the unsuccessful helpers dying afterwards. After *k* helpers have been recruited to the local patch, the cycle then repeats.

### 2.2 Fitness

The expected number *w_ij_* of offspring who successfully establish themselves in a patch with *i* helpers and born from a mutant adult breeder in a patch with a total number of *j* helpers is then given by

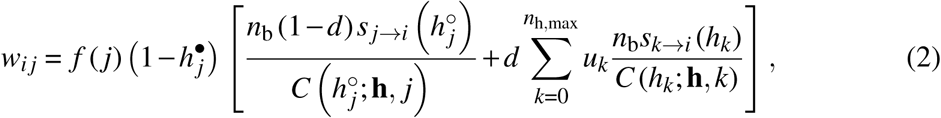

where *f*(*j*) reflects the total number of surviving newborns produced by the focal adult breeder, a proportion 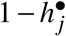 of which develop as juvenile reproductives (rather than helpers). These juvenile reproductives then go on to compete for any of the *n*_b_ available breeding positions in the natal patch with probability 1 − *d* (first part in straight brackets), or in a random, remote patch with probability *d* (second part in straight brackets), where *u_k_* reflects the population-wide frequency of patches currently containing 0 ≤ *k* ≤ *n*_h,max_ helpers. Philopatric reproductives compete with a total number of 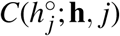 philopatric and immigrant offspring (see eq. [S1] in the Online Supplement), which is a function of (i) the average tendency 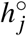 expressed by any locally born newborn to develop as a helper, (ii) the population wide tendencies **h** = [*h*_0_, *h*_1_, …, *h*_*n*_h,max__] to become helpers in any remote patch and (iii) the current number of helpers *j* in the local patch. Finally, after successful establishment, the probability that the newly established breeder is accompanied by *i* helpers in the next generation is then given by 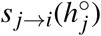 (see eq. 1). The expected number of offspring who successfully compete in remote patch can then be derived in a similar fashion.

#### 2.3 Evolutionary dynamics

We use a direct fitness method (also called neighbour-modulated fitness [32, 33]) to calculate evolutionary change 𝓗_*k*_ in the tendency to help when born on a patch containing *k* helpers.

According to a standard result [34–36], 𝓗_*k*_ is then given by

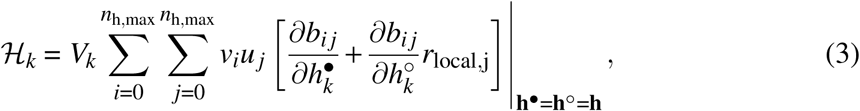

where *V_k_* is a term that is proportional to the amount of additive genetic variance in the helping tendency *h_k_*. Next, *v_i_* and *u_j_* are the individual reproductive values and stable class frequencies of adult breeders which are in a patch with *i* helpers, which are obtained from the dominant left and right eigenvectors of the resident transition matrix (see eq. [S3]). Finally, the relatedness coefficient *r*_local,*j*_ reflects the relatedness between a focal adult breeder and any distinct patch mate when being helped by *j* helpers (see eq. [S4]). As we have not been able to find analytical solutions to find *h_k_*, we developed an algorithm in C++ (source code available at https://doi.org/10.5281/zenodo.1421729) to numerically find the convergence stable values of the helping tendencies **h** (see Online Supplement). Throughout, we assume that helper-dependent fecundity of a focal breeder in a patch with *j* helpers is given by the function *f*(*j*) = (1/*n*_b_)(*a* + *mj^z^*), where *a* is the baseline productivity of a patch without helpers, *m* is the strength with which productivity increases with increasing helper number and *z* reflects whether the increase happens in linear accelerating or decelerating fashion. We assume that the benefits of helping are equally shared among all *n_b_* breeders.

#### 2.4 Results

##### 2.4.1 Result 1: individuals who receive more help evolve higher tendencies to help themselves

To assess how the presence of helpers in early life affects an individual’s tendency to help others, we focus on a scenario were maximally *n*_h,max_ = 5 helpers can be recruited to a local patch and where helper number has a linear effect on local productivity. Results are, however, robust to different values of *n*_h,max_ (Figure S1) or cases where helper numbers increase local productivity in a decelerating fashion (Figure S2).

Figure 1A shows that early-life effects on the development of adult helping behaviours are adaptive, as the probability of helping in adulthood is strongly dependent on the amount of help received in early life (as measured by the number of helpers in the local patch). Moreover, we find that those individuals who have experienced an intermediate number of helpers at birth (e.g., *n*_h_ = 1,2,3) are most likely to develop as helpers themselves in later life, whereas adults who have experienced either a very large amount of help (e.g., *n*_h_ = 5), and particularly those who have received no help at all (*n*_h_ = 0) are less likely to become helpers themselves. Finally, we find that early-life effects also enhance the evolutionary scope for helping, relative to populations which are constrained from using early-life information (unconditional helping): for example, for high rates of dispersal (*d* = 0.6), helping only evolves when individuals can use early-life information but not for populations where helping is un-conditional (compare solid vs dotted green lines in Figure 1A).

**Figure 1.**
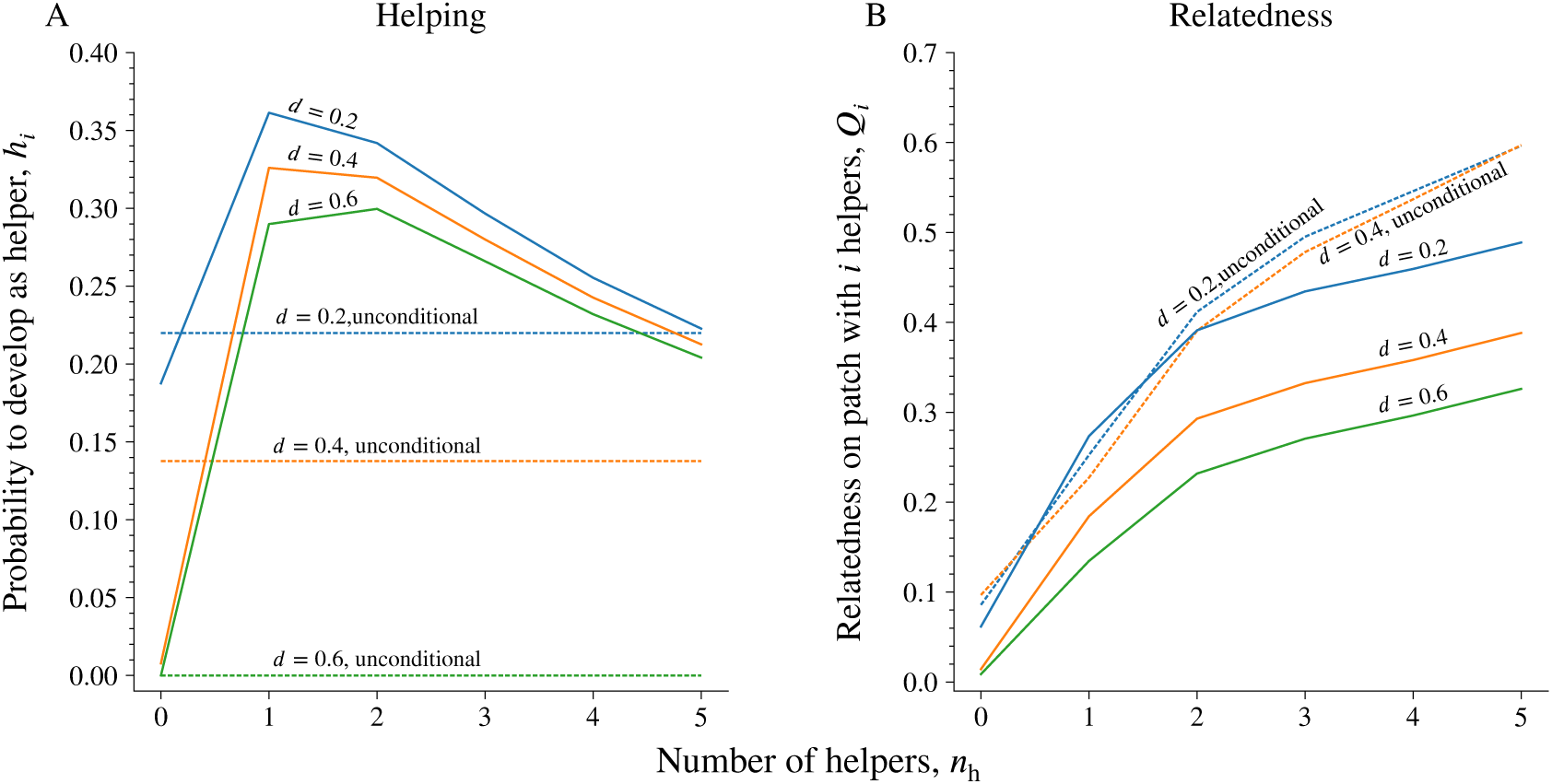
The evolution of developmentally plastic and unconditional helping behaviours in patches that contain 0 ≤ *n*_h_ ≤ 5 helpers in early life, for three different values of juvenile dispersal *d* (panel A). When helping is developmentally plastic, individuals develop higher levels of help in patches where helpers are present in early life (n_h_ > 0) relative to patches where help in early life is absent (*n*_h_ = 0). Panel B depicts the corresponding relatedness coefficients, showing that relatedness is higher in patches with more helpers, as these are more productive (hence making it more likely that philopatric offspring claim breeding spots). Note that when *d* = 0.6, unconditional helping does not evolve, hence we only have *n*_h_ = 0. The corresponding line in panel A is hence only drawn for the purpose of illustration, while we do not depict a corresponding relatedness coefficient for *d* = 0.6 and unconditional helping. See also Figure S1 for *n*_h,max_ ∈ {3,10,20} and Figure S2 when productivity increases in a decelerating (rather than linear) fashion with increasing numbers of helpers. Parameters: *n*_b_ = 2, *z* =1.0, *n*_h,max_ = 5, *m* = 5, *g* =1.0.

To understand the evolution of early-life effects on helping, Figure 1B depicts the coefficient of consanguinity [32, 37] between two distinct breeders for patches with different numbers of helpers. We find that, once helping evolves, relatedness is highest in those patches which contain the largest numbers of helpers and lowest in patches where helpers are absent. This is because a larger number of helpers increases the fecundity of the local group, so that any vacant breeding spots are more likely to be claimed by locally born juveniles (rather than by remotely born offspring). Consequently, helpers and breeders are more likely to be related in the next generation, thus favouring stronger helping tendencies in patches that currently contain high numbers of helpers (note that relatedness varies even more strongly with helper number when helping is unconditional [dotted lines in Figure 1B], suggesting that any mechanism which would make helping sensitive to early-life experiences of numbers of helpers should rapidly invade in populations with unconditional helping).

Due to the feedback between developmentally plastic helping and relatedness, we would thus expect an outcome where experiencing a high (or a low) number of helpers during childhood makes individuals more likely (or less likely) to perform helping behaviours themselves as adults, suggesting a positive feedback between early life and later life sociality. At the same time, however, high productivities of patches with large numbers of helpers result in a rapid saturation of the available number of helper vacancies, which explains why the tendency to help is only maximized on patches with an intermediate number of helpers (see Figure 1A). However, when saturation of helper positions is less likely (e.g., because of a higher number of helper vacancies), helping tendencies are often maximized for patches with a larger intermediate number of helpers (e.g., see Figure S1), while also the overall helping tendencies attain higher values.

##### 2.4.2 Result 2: Helper presence as a predictor of a social future

We then quantified the longer-term consequences that result from the presence (or absence) of helpers. When helper development depends on the current amount of help received (Figure 2A), we find that even for relatively high levels of dispersal (i.e., *d* ≈ 0.6) the current number of helpers is highly predictive of the number of helpers in the future (see also the autocorrelations in Figure 3, blue line). For example, patches that currently have no helpers (*n*_h_(*t*) = 0) are extremely unlikely to recruit any helpers in the future. Similarly, patches that currently have the maximum number of helpers (e.g., *n*_h_(*t*) = 5, thick blue lines) are highly likely to have a large number of helpers again in the future.

**Figure 2.**
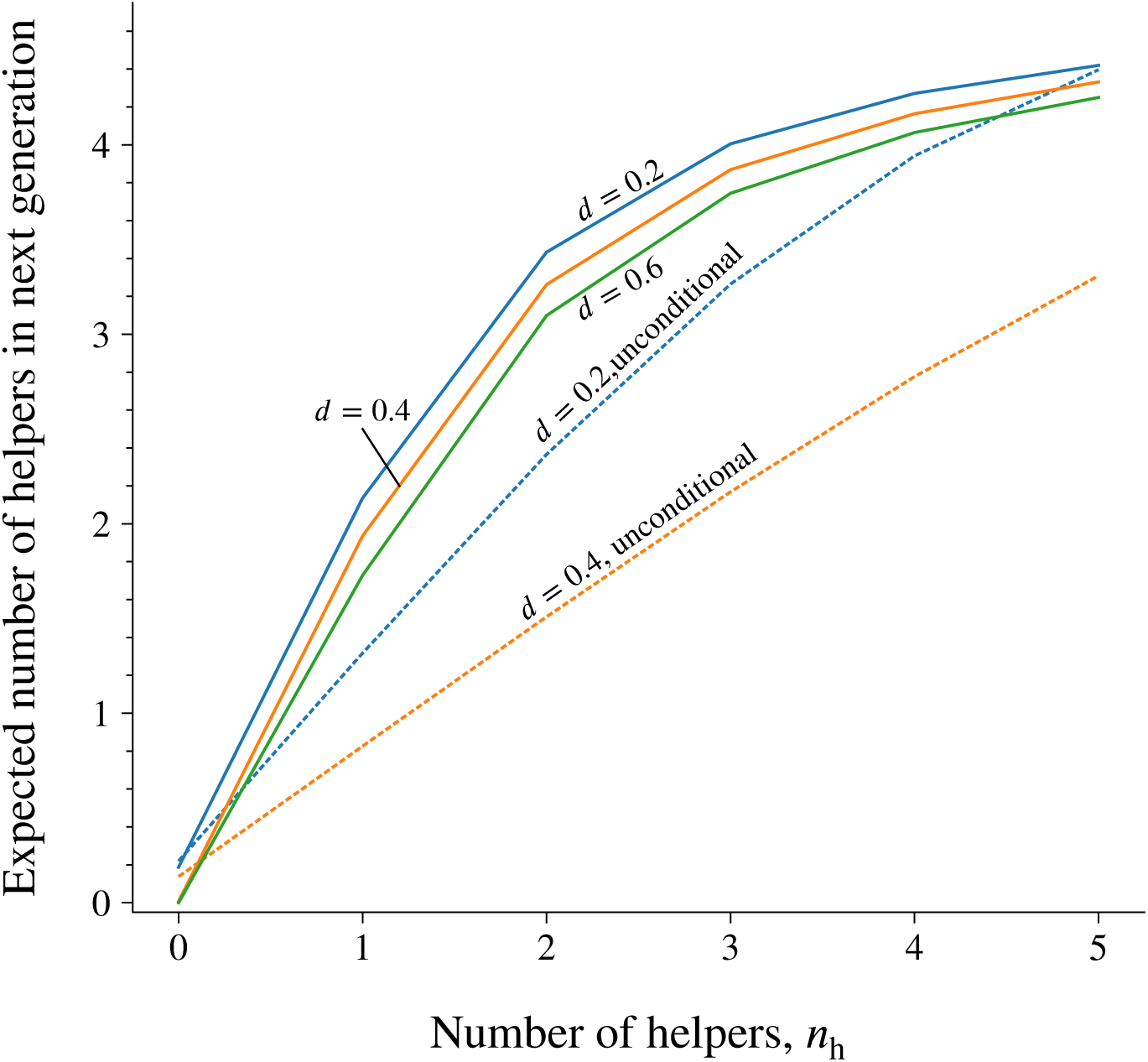
The expected number of helpers recruited in the next timestep increases with current number of helpers in the local patch. It does so much more rapidly in case helping is developmentally plastic (solid lines). Note that when *d* = 0.6, unconditional helping does not evolve, hence the expected number of helpers is always equal to 0. Parameters: *n*_b_ = 2, *z* = 1.0, *n*_h,max_ = 5, *m* = 5, *g* = 1.0.

**Figure 3.**
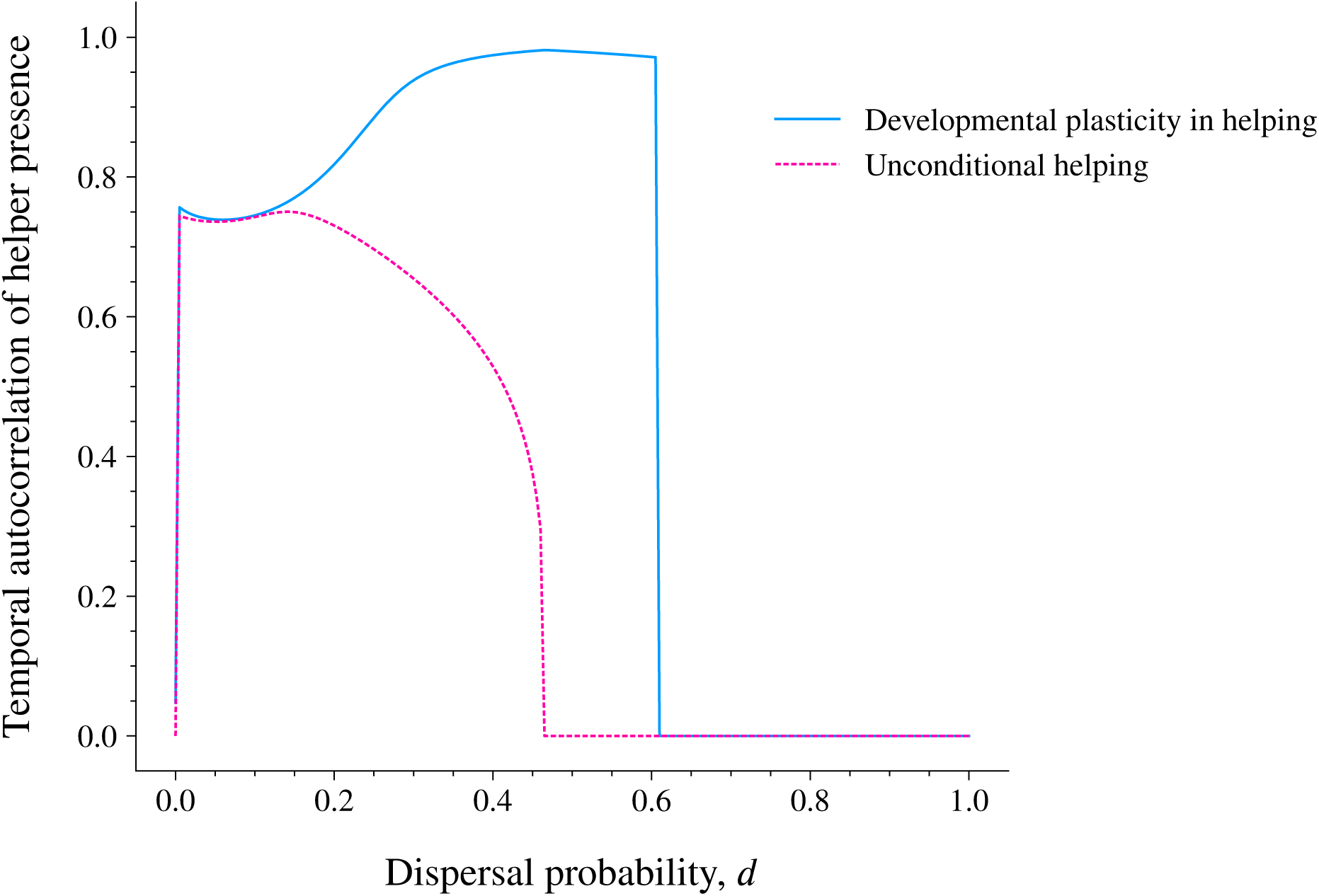
Temporal autocorrelation in patterns of presence versus absence of helpers between parental and offspring generations, for populations in which help is developmentally plastic (solid blue line) versus unconditional (dotted pink lines), while varying the probability of juvenile dispersal *d*. Parameters: *n*_b_ = 2, *z* = 1.0, *n*_h,max_ = 5, *m* = 5, *g* = 1.0.

By contrast, when helping is unconditional, help only evolves when dispersal is more limited (i.e., *d* ≈ 0.45), but even then the number of helpers experienced in the current generation is a much poorer predictor of the amount of help received in the future (see also Figure 3). Only in populations where dispersal becomes strongly limited, relatedness is high overall, so that helping evolves regardless of the current number of helpers in the local patch. Indeed, when *d* is small, the autocorrelation in the presence of helpers is identical for populations where help is either unconditional or conditional on the number of helpers experienced at birth.

## 3 Discussion

Here, we have shown that when individuals are free to modulate their social behaviour as adults (specifically their tendency to become non-reproductive helpers) in response to the social environment they experience in early life, selection favours positive social sensitivity. Greater experience of prosocial behaviour in early life then results in a greater tendency to behave prosocially during adulthood, while a reduction in prosocial behaviour is an adaptive response to social adversity in early-life, potentially explaining empirical patterns in humans [10] and other societies (e.g., [4]).

Our model predicts a positive relationship between early-life social experience and later-life social behaviour because helping promotes local productivity, thereby increasing local relatedness, and greater local relatedness in turn favours more helping. To understand this, focus on a local group with many helpers: this group will produce a large number of offspring, hence increasing the probability that a local breeding spot will be claimed by a locally born (rather than a remotely born) offspring in the next generation. In turn, this results in an increase in local relatedness (see Figure 1B), favouring a high tendency to develop as a helper. By contrast, patches which currently contain few helpers are less productive, ultimately resulting in a lower relatedness and a lower tendency to help. Of course, this kind of feedback between sociality and relatedness will only develop when individuals can adjust prosocial behaviour in response to juvenile cues that are predictive of local relatedness experienced as an adult. Our model shows that the experience of being helped in early life can serve as a reliable cue of expected relatedness in this way, thus driving developmental plasticity in later-life social behaviour.

Once positive social sensitivity has evolved, the intergenerational propagation of prosocial behaviour itself amplifies the benefits of helping, because an individual who becomes a helper not only boosts the fecundity of related breeders in the current generation, but also increases the tendency to help among progeny that remain on the local patch. Helping, in other words, ends up providing longer-term as well as shorter-term benefits. In a series of seminal models, Lehmann [38, 39] has previously shown that persistent benefits, which impact on the fitness of later generations, are particularly favourable for the evolution of helping, because they provide a partial escape from the constraints of local kin competition. These models, however, start from the assumption that the benefits of helping behaviour persist over time, as seems likely to be true for many beneficial modifications of the local environment such as construction or maintenance of a nest or burrow. Our model shows that even if helping has no such physically persistent effects, and only boosts the fecundity of breeders in the current generation, it may nevertheless end up yielding longer-term benefits because of the inter-generational propagation of prosocial tendencies. Consequently, we argue that long-term benefits of helping may be a much more common feature of kin selection than hitherto appreciated, and can be interpreted as a form of social niche construction (by analogy with the physical niche construction emphasised by Lehmann).

Previous models of the evolution of early life effects have focused chiefly on adaptation to fluctuations in the abiotic environment (e.g., [13, 15, 18, 40–43]) with surprisingly little attention given to social sensitivity (as previously noted in [3]). A key prediction of existing theory is that environmental conditions need to be sufficiently autocorrelated with later-life environmental conditions. However, some studies suggest that autocorrelations from climatic timeseries are, in fact, small and thus cannot readily account for the widespread occurrence of early-life effects (e.g., [44, 45]). Our model, however, shows that variation in the social environment can drive the evolution of early-life effects, even in the absence of autocorrelations in the abiotic environment, because social sensitivity itself generates high autocorrelations between parental and offspring social environments (see Figure 3). Hence, our study suggests that the social environment may in general play a more important role in the evolution of early-life effects than does the abiotic environment (see also [46]).

Our model suggests a number of possible directions for future work: as discussed above, one key prediction is that increased levels of social behaviour result in increases in relatedness, thus creating a positive feedback loop. A typical consequence of positive feedback loops is that they often result in alternatively stable states [47]. Indeed, Figure 2A suggests that developmental plasticity may well result in a social polymorphism at the patch level, in which a highly prosocial state is maintained persistently in some subpopulations, while others become locked into persistently less prosocial states (note that neither state will persist indefinitely, due to demographic stochasticity in recruitment of helpers to a patch, but that positive trans-generational feedback on social behaviour will tend to maintain differences between patches for longer than would otherwise be the case). Such persistent polymorphism represents a group-level analog of the persistent differences in individual behaviour that emerge during early life and are maintained in models of personality evolution [48, 49]; one might even speak of the emergence of ‘collective personalities’. However, a complete analysis of these consequences of developmental plasticity in helping is beyond the scope of the current paper and would merit further study.

Another potential subject for future research is to investigate whether a positive feedback between social behaviour and relatedness is indeed a general outcome, or whether this depends on specific model assumptions. For instance, one assumption that deserves further scrutiny is that helping in our model affects fecundity, rather than survival [50]. We would predict that when increasing numbers of helpers instead boost breeder survival (rather than fecundity) [50, 51], offspring should still evolve to provide more help as adults if they receive more help in early life. This is because under these circumstances more prosocial patches contain breeders who live longer, which are therefore more likely to be related to the helpers in the local patch. However, the prediction that survival, rather than fecundity selection, again favours positive social sensitivity remains to be verified.

A potentially interesting scenario in which our finding of a positive feedback might be more dramatically affected is when increased levels of helping result in disproportionally more dispersing juveniles, hence resulting in a negative feedback between helping and relatedness. Such a scenario may apply when dispersal is energetically demanding and hence only possible when a minimum level of help is provided to offspring. In this case, patches with few (many) helpers produce more (fewer) philopatric offspring, resulting in increased (decreased) levels of relatedness in the next generation. We would predict that early-life sensitivity in social traits might still evolve in this context, but in an opposite fashion, such that early-life social adversity favours more (rather than less) prosociality later in life. Studies which set out to assess when positive versus negative feedbacks between sociality and relatedness are to be expected would be of great interest. Overall, our model shows that early-life effects in social contexts can be adaptive, but highlights the need for further study to understand their ecological significance.

## Acknowledgements

BK is funded by a Leverhulme Trust Early Career Research Fellowship (ECF 2015-273). Ana Duarte is thanked for discussions of early-life effects. We thank the University of Exeter for providing computational resources through the Carson computing cluster.

